# A mosaic bulk-solvent model improves density maps and the fit between model and data

**DOI:** 10.1101/2021.12.09.471976

**Authors:** Pavel V. Afonine, Paul D. Adams, Oleg V. Sobolev, Alexandre Urzhumtsev

## Abstract

Bulk solvent is a major component of bio-macromolecular crystals and therefore contributes significantly to diffraction intensities. Accurate modeling of the bulk-solvent region has been recognized as important for many crystallographic calculations, from computing of *R*-factors and density maps to model building and refinement. Owing to its simplicity and computational and modeling power, the flat (mask-based) bulk-solvent model introduced by Jiang & Brunger (1994) is used by most modern crystallographic software packages to account for disordered solvent. In this manuscript we describe further developments of the mask-based model that improves the fit between the model and the data and aids in map interpretation. The new algorithm, here referred to as *mosaic bulk-solvent model*, considers solvent variation across the unit cell. The mosaic model is implemented in the computational crystallography toolbox and can be used in *Phenix* in most contexts where accounting for bulk-solvent is required. It has been optimized and validated using a sufficiently large subset of the Protein Data Bank entries that have crystallographic data available.

**Synopsis:** A mosaic bulk-solvent method models disordered solvent more accurately than current flat bulk solvent model. This improves the fit between the model and the data, improves map quality and allows for the solution of problems previously inaccessible.

## 1. Introduction

Bulk solvent (or disordered solvent) occupies up to 90% of crystallographic unit cell volume of macromolecular crystals and contributes significantly to the diffraction data, mostly to low-resolution reflections (Weichenberger *et al*., 2015; Urzhumtsev & Lunin, 2019). It is therefore important to include its contribution into the model-calculated structure factors to ensure the best map quality and successful refinement of the atomic model. A flat (or mask-based) bulk solvent model (Jiang & Brünger, 1994) is the option of choice in most crystallographic software packages such as *CNS* (Brunger *et al*., 1998), *Refmac* (Murshudov *et al*., 2011) and *Phenix* (Liebschner *et al*., 2019). This model is simple while still modeling the solvent contribution well, and typically includes the following steps:

1. given an atomic model, calculate a set of structure factors **F**_*calc*_(**s**) that corresponds to the set of experimental amplitudes *F*_*obs*_(**s**);
2. define a solvent mask in the unit cell that is a binary function calculated on a grid with zero inside the molecular region and one outside;
3. compute the Fourier transform of this mask to obtain Fourier coefficients **F**_*mask*_(**s**) with Miller indices matching *F*_*obs*_(**s**);
4. Define the total model structure factors as

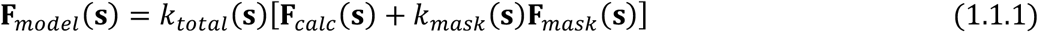

and compute resolution-dependent scale factors *k*_*total*_(**s**) and *k*_*mask*_(**s**) such that **F**_*model*_(**s**) fits the measured amplitudes *F*_*obs*_(**s**) as well as possible (Jiang & Brünger, 1994; Fokine & Urzhumtsev, 2002; Afonine *et al*., 2005; Fenn *et al*., 2010; Afonine *et al*., 2013). This flat bulk-solvent model assumes exactly the same solvent density everywhere in the unit cell outside the atomic model. This makes it rather straightforward to envision where and why this model may be suboptimal.

*First*, the molecule shape and the crystal packing may allow for several isolated regions that can vary in size (Fig.1; Appendix A) and can potentially be filled at different occupancy rates, including completely empty regions (Liu *et al*., 2008; Matthews & Liu, 2009). Assuming exactly the same occupancy in all these regions can result in noisy difference maps and suboptimal model-to-data fit.

**Figure 1.**
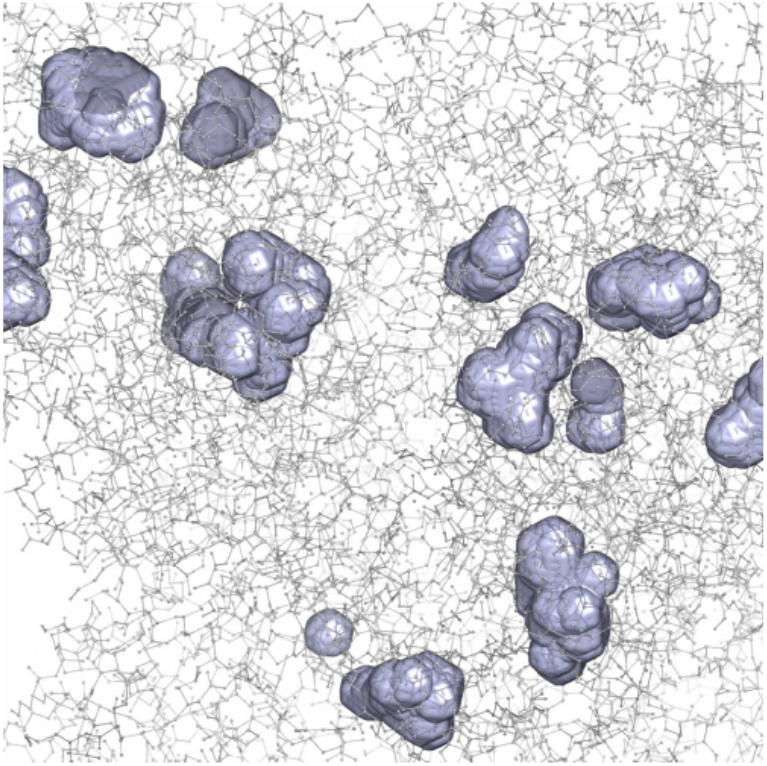
Example of isolated regions (blue) inside macromolecule (PDB code 1bxs).

*Second*, the flat solvent model is unaware of alternative conformations. For example, if a residue side chain occupies two locations, then bulk-solvent will be fully excluded from both locations. This is likely to result in patches of residual density artifacts overlaying each of the conformers, or these artifacts will be minimized by incorrectly refined occupancies and/or B-factors.

*Third*, a solvent region may contain different chemical entities. For example, there may be parts systematically occupied by disordered components such as lipids both in isolated regions (Lunin *et al*., 2001) or as a part of the main solvent region (Sonntag *et al*., 2011) This requires different bulk solvent parameters for the different solvent components.

*Forth*, some isolated regions may actually be occupied by poorly ordered ligands. Applying a solvent model inside these regions may flatten out the poor ligand density further and impede its interpretation (Liebschner *et al*., 2017).

Previously, some improvements of the bulk-solvent model have been reported. They mostly concerned improvement of scale factors (Fokine & Urzhumtsev, 2002; Afonine *et al*., 2005; Afonine *et al*., 2013) and the treatment of solvent at the macromolecule-solvent interface (Fenn *et al*., 2010). Further improvements are required to address the limitations of the current model outlined above.

Here we introduce a new bulk-solvent model that we refer to as *mosaic bulk-solvent model*. It assumes bulk-solvent can vary across the unit cell volume and be present at not necessarily full occupancy. Individually modelling the mean solvent density in distinct regions is expected to improve the first three situations described above, however its straightforward application may complicate ligand identification. In this work we address the first situation above, and intend to address the other three in future work.

## 2. Method: mosaic bulk solvent

### 2.1. Mosaic model

#### 2.1.1. Generalities

The classic bulk-solvent modeling implicitly supposes that the whole region outside the macromolecule, called the bulk solvent region, is uniformly filled by the same solvent. This may not reflect the reality as discussed in the Introduction. The mosaic model treats the solvent region as an accumulation of several non-intersecting regions, *n* = 1, …, *N* (Appendix A) where the mean solvent density may eventually vary across the regions. We define each region by its mask Ω_*n*_(**r**), *n* = 1, …, *N*, where Ω_*n*_(**r**) is equal to 1 inside the region *n* and 0 outside it. In what follows we consider Ω_0_(**r**) to be the macromolecular mask. These non-intersecting regions, including the macromolecular region, cover the whole unit cell, which means

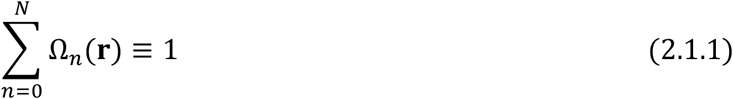

for any point **r**. The total model structure factor is then defined as a sum of contributions from all these regions:

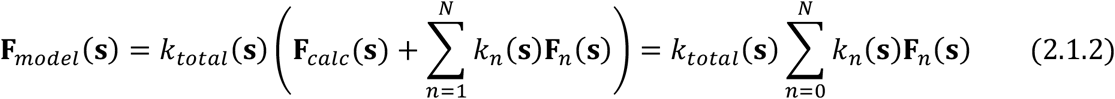

where **F**_0_(**s**) = **F**_*calc*_(**s**), *k*_*total*_(**s**) and *k*_*n*_(**s**), *n* = 1, …, *N*, are unknown scale factors that can be obtained by fitting of (2.1.2) to the experimental structure factor amplitudes *F*_*obs*_(**s**) in resolution shells, *i* = 1, …, *I*. This is a generalization of the previous *Phenix* model that considers a single bulk solvent region, *N* = 1 (Afonine *et al*., 2013; in what follows referred to as A-13). The new procedure is based on several assumptions:

-*k*_*total*_(**s**) falls off with the resolution and can be approximated by a Gaussian;
-*k*_*total*_(**s**) is anisotropic and obeys the crystal symmetry;
-All scale coefficients *k*_*n*_(**s**) are step functions of the resolution; this means *k*_*n*_(**s**) are assumed constants within each resolution shell and therefore resolution shells need to be sufficiently thin for this assumption to be reasonable;
-For the coefficients *k*_*n*_(**s**) we do not assume any particular analytic form (Urzhumtsev & Podjarny, 1995; A-13);
-In protein crystallography, *k*_*mask*_(**s**) is non-negative and usually varies around 0.3 eÅ^-3^ reflecting the mean electron density of the solvent (Fokine & Urzhumtsev, 2002);
-The contribution of bulk solvent vanishes at approximately 3-4 Å resolution and higher;
-Regions of bulk solvent should have a volume sufficient to accommodate a few solvent molecules.

In this model, each solvent component has its own scattering function and coefficients *k*_*n*_(**s**) are independent from each other.

A simplified variant of the mosaic model can be envisaged by assuming that all regions contain exactly the same kind of solvent, with exactly the same scattering properties, and the only difference is the occupancy of these regions. This means that the bulk solvent structure factors are defined as

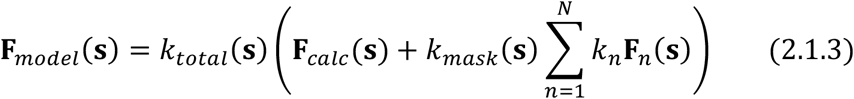

An advantage of this model is that it only uses a single parameter *k*_*n*_ per region, and the total number of independent parameters reduces from (*N* + 1)*I* to (*N* + 1) + *I*, where *I* is the number of resolution shells. Unlike (2.1.2) here the coefficients *k*_*n*_ are resolution independent and can be obtained by one of the algorithms described below using all reflection data at once. The regions with a low occupancy are candidates for removal, while regions with high *k*_*n*_ are likely to indicate ordered or semi-ordered solvent or solvent with scattering properties being different from the rest.

To simplify expressions, in what following we omit the argument (**s**) and the index *i* of the resolution shell when this does not lead to ambiguities.

#### 2.1.2 Model parameters

The bulk solvent contribution to the total model structure factor (2.1.2) is defined by the shape of the mask and the grid on which this mask is calculated according to Jiang & Brünger (1994); in what follows we refer to as JB-94. The shape is defined by two radii, *r*_*shrink*_ and *r*_*solv*_ (called *r*_*probe*_ in JB-94). The grid step is usually taken as *d*_*min*_/4 (JB-94) while Rees *et al*. (2005) argued for *d*_*min*_/5 or even *d*_*min*_/10 for low-resolution projects. In fact, we show that the grid step for the calculation of the bulk solvent mask is not related to the resolution of the data and can instead be defined as a universal value for all structures (Appendix B). The universal and highly correlated values of the two radii can be chosen given the grid step used. We find *r*_*solv*_ = 1.1 Å, *r*_*shrink*_ = 0.9 Å and *d*_*step*_ = 0.6 Å as the optimal choice for these parameters (Appendix B).

#### 2.1.3. Bulk solvent masks

The solvent region defined above may consist of several connected components. Connectivity analysis (Lunina *et al*., 2003) is used to identify them while also considering the periodicity of the crystal. A region will have symmetry-related copies when the crystal symmetry is different from primitive triclinic (P1). We refer to “the coefficients from the region *n*” considering that all symmetry-related copied of this region have been taken together.

The molecular, and therefore the solvent mask(s), depends on the atoms of the atomic model. We expected that including hydrogen atoms for the mask calculation step may improve the model; however, the result was just the opposite (Appendix B). Therefore, hydrogen atoms were ignored in the mask calculation.

The bulk solvent region may contain some points that physically cannot be occupied by solvent molecules, e.g., regions that are too small to accommodate a solvent molecule or isolated grid nodes or points that form lines or two-dimensional surfaces. The mosaic procedure excludes these regions from (2.1.1) by applying volume cutoffs and other map analysis algorithms (Appendix C).

### 2.2. The protocol

The procedure starts from the atomic model and the set of observed structure factor amplitudes, *F*_*obs*_(**s**). The atomic model is used to calculate the bulk solvent mask as described in JB-94 using optimal parameters for the grid step and radii (§ 2.1.2). Then connectivity analysis is applied to this mask to find all isolated regions in it. Masks that survive the region filtering criteria (Appendix C) are then used to calculate corresponding structure factors to be used in (2.1.2). We note that this is not the only way to divide the bulk solvent mask into regions. For example, mask regions can be partitioned on the basis of disordered lipid belts in membrane proteins (Sonntag *et al*., 2011); not discussed in this work.

To estimate the initial values of *k*_*total*_(**s**) the A-13 calculation is performed using the largest mask region. Total model structure factors **F**_*model*_(**s**) obtained here are used to calculate a difference map. This map can be thought of as a residual OMIT map (Bhat, 1988, and references therein) for the smaller individual solvent regions and thus can be used for additional region filtering as described in Appendix C. Also, the phases of **F**_*model*_(**s**) are used as initial values for iterative phased search of scales *k*_*n*_(**s**) (§2.4.4; Algorithm 4).

Observed amplitudes *F*_*obs*_(**s**) or intensities *I*_*obs*_(**s**) and scaled 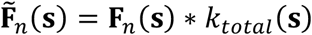 are the inputs to one of the protocols to calculate *k*_*n*_(**s**) coefficients in (2.1.2). Calculation of *k*_*n*_(**s**) are performed in resolution shells as described below. The procedure is repeated iteratively until convergence with *k*_*total*_(**s**) and *k*_*n*_(**s**) being updated at each iteration. Typically, 2-3 iterations are required to reach convergence. Values for *k*_*n*_(**s**) are checked at each iteration: if *k*_*n*_(**s**) for a particular region is unrealistic (for example, negative) the corresponding **F**_*n*_(**s**) is permanently removed from (2.1.2). Once converged, the **F**_*model*_(**s**) can be calculated (2.1.2) and used as needed.

### 2.3. Resolution shells

The coefficients *k*_*n*_(**s**) are calculated as constants in resolution shells. The choice of resolution shells is similar to that used in A-13 with two differences. First, previous experience (A-13) shows that coefficients *k*_*n*_(**s**) vanish at approximately 3 Å resolution and higher, therefore these reflections are excluded from calculations. Second, A-13 used approximately *N*_*refl*_ = 100 reflections for the lowest resolution shell and a uniform logarithmic scale for the following shells (Urzhumtsev *et al*., 2009). In the mosaic model, there are *N* independent regions and *N* independent coefficients *k*_*n*_(**s**) per resolution shell instead of one. To keep the same data-to-parameters ratio as in A-13, we merge adjacent bins so that each new merged bin has the required number of reflections. Usually, only a few lowest-resolution shells are concerned since the number of reflections per shell increases rapidly with resolution (Table 1 in A-13).

### 2.4. Algorithms to calculate coefficients in (2.1.2)-(2.1.3)

#### 2.4.1. Algorithm 1: sequential search

Algorithm 1 is essentially the previous algorithm A-13 applied iteratively to the connected regions one after another, *n* = 1, …, *N*, keeping the contribution from other regions fixed. This algorithm is sensitive to the initial values of the scale factors; poor initial estimates for some of them may result in incorrect estimates for the others and the failure of the whole procedure. Also, the algorithm is computationally expensive and thus can’t be readily applied when the number of regions is large (Appendix A).

#### 2.4.2. Algorithm 2: iterative one-step search

This algorithm searches simultaneously for all values of the coefficients *k*_*n*_ in each resolution shell *i*, where this coefficient is constant, minimizing the residual

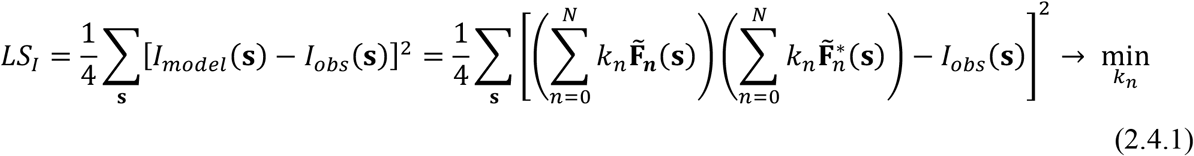

Here the sums are calculated over the reflections of the given shell (we omit its index *i*). This expression can be rewritten as

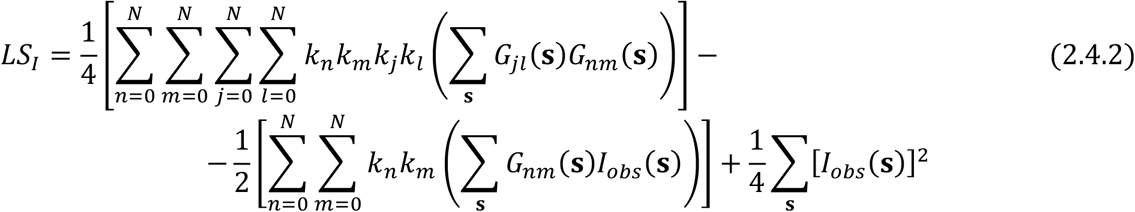

where

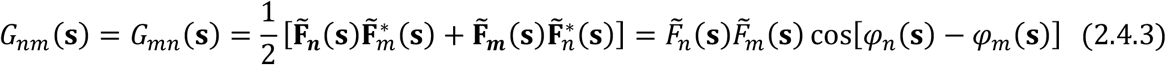

The polynomial of the fourth degree (2.4.2) with respect to individual scale factors *k*_*n*_ can be minimized using a standard approach, e.g. L-BFGS (Liu & Nocedal, 1989). The derivatives of *LS*_*I*_ with respect to *k*_*j*_, *j* = 0, … *N*, required by the minimizer, are

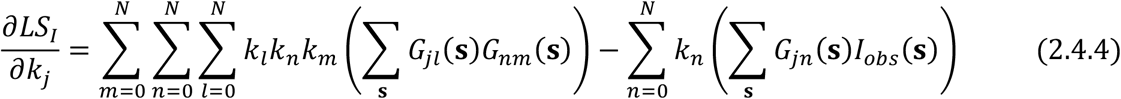

#### 2.4.3. Algorithm 3: non-iterative two-steps search

In this algorithm, instead of using iterative minimization methods, we search for the minimum of (2.4.2) analytically, requiring no start values. First, we introduce (*N*+1)^2^ intermediate parameters

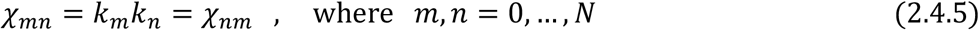

We search first for their values that we decompose later into individual coefficients *k*_*n*_. The function (2.4.2) becomes a quadratic function of these new variables

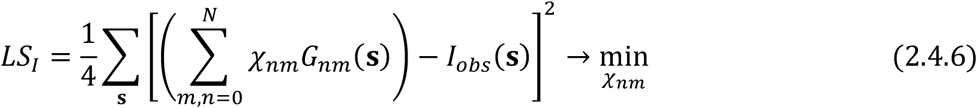

The minimum of *LS*_*I*_ can be found as a solution of a system of linear equations with respect to its parameters. After excluding the redundant variables, we stay with ½(*N*+1)(*N*+2) equations for the independent variables *χ*_*min*_, 0 ≤ *m* ≤ *n* ≤ *N* :

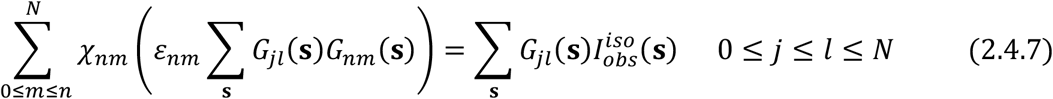

Here *ε*_*mn*_ = 1 if *m* = *n* and *ε*_*mn*_ = 2 otherwise. This system can be solved by standard approaches. In the second step, we search for *N*+1 scale coefficients *k*_*n*_ minimizing

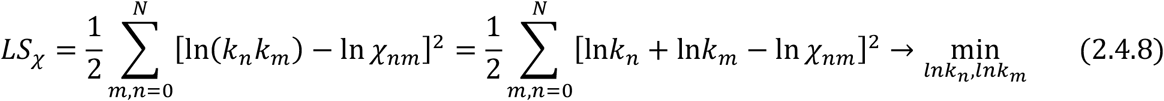

The minimum of this function, quadratic with respect to ln*k*_*n*_, can be found analytically as a solution of the system of linear equations

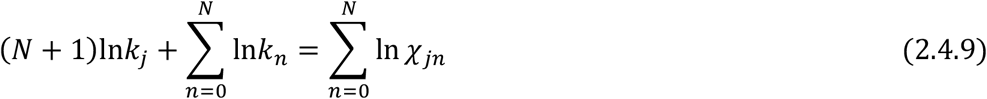

giving

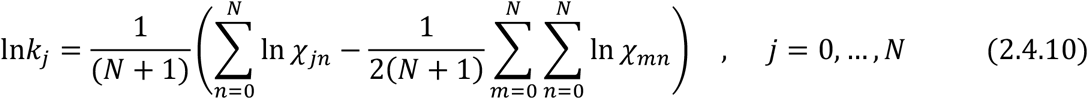

Finally, we recover *k*_*n*_ from ln*k*_*n*_.

While this algorithm requires neither iterations nor initial values of the scale factors, its serious disadvantage is the large dimension of the system of equations (2.4.7), a need to use a square matrix of the dimension 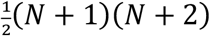, and sensitivity to rounding errors. It works well for synthetic numerical test examples but is impractical when applying to real structures. We describe it here for the sake of completeness but do not use it in the tests below.

#### 2.4.4. Algorithm 4: iterative phased search

With this algorithm, we try to avoid both an iterative minimization of a function of many variables (Algorithm 2) and use of a large system of equations (Algorithm 3). To do so, instead of comparison of intensities, we compare structure factors as complex values. The phase values *φ*_*obs*_(**s**) can be taken as those of the model structure factors (2.1.2)

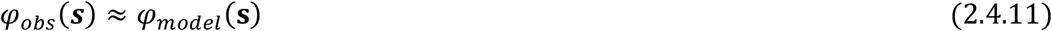

which is a reasonable assumption for a nearly finalized model (the scenario when mosaic model is ought to be used).

We express the best fit of the complex structure factors as

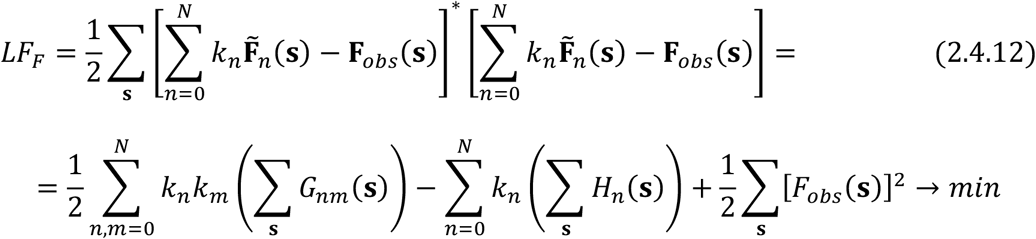

where *G*_*mn*_(**s**) are defined previously by (2.4.3) and *H*_*n*_(**s**) are defined similarly as

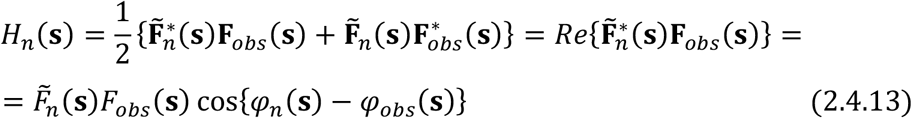

The condition (2.4.12) results in a system of *N*+1 linear equations with respect to *k*_*n*_:

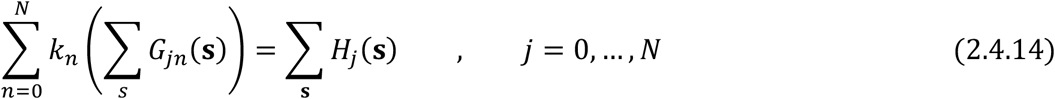

Several iterations may be required solving (2.4.14), improving model structure factors (2.1.12) and respective phase values (2.4.11) and updating *H*_*n*_(**s**) (2.4.13).

### 2.5. Mosaic model and ordered ligands

In the algorithms above, we suppose that all isolated regions have disordered content. However, some regions may be occupied by structured ligands for which an atomic model can be proposed. Filling these regions with the mean solvent density by A-13 or with a specifically adjusted density by the mosaic model is likely to reduce the contrast of the ligand or even obliterate it completely in residual maps (Liebschner *et al*. (2017). Figure 2 shows two examples of ligands in small isolated regions. In the absence of an atomic ligand model these regions are filled with the bulk-solvent (Fig. 2a). Omitting bulk solvent from these regions shows clear and unambiguous ligand density (Fig. 2b). Applying A-13 with the default mask damages the residual map (Fig. 2c) and, expectedly, applying mosaic model deteriorates the map even more (Fig. 2d), which is undesirable.

**Figure 2.**
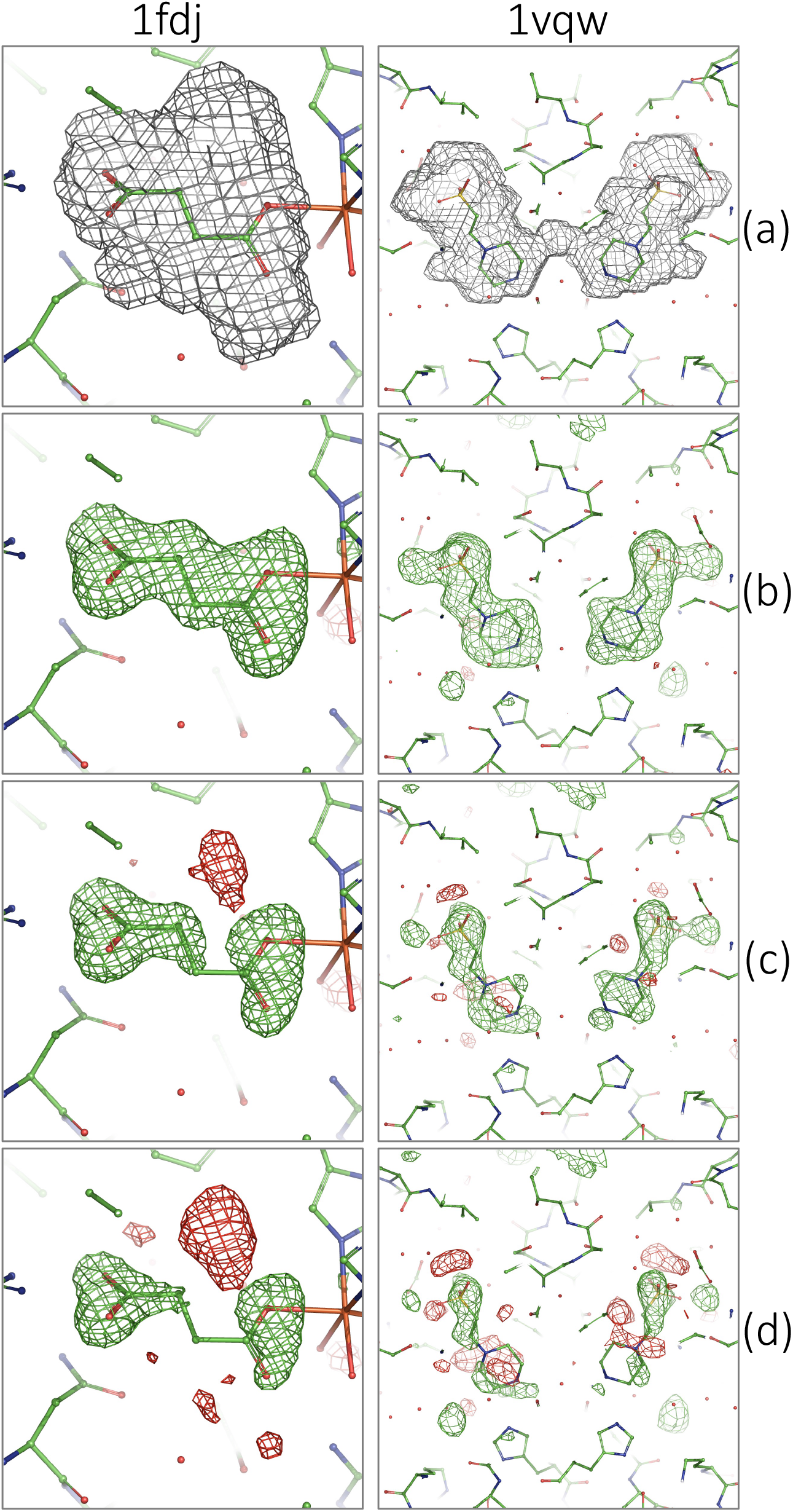
Two ligands in isolated region (PDB codes: 1fdj and 1vqw). (a) Bulk-solvent mask. mF_obs_-DF_model_ ligand OMIT maps with F_model_ calculated: (b) using one largest bulk-solvent mask, (c) as in A-13 and (d) using mosaic bulk solvent model. Maps (b-d) are contoured at ±3*σ*, green – postive, red – negative.

The solution that we propose is to calculate and simultaneously use two residual maps. One map is calculated using the current complete model and mosaic approach. The second map is calculated using only the main component of the bulk-solvent mask, i.e., the mask with all other isolated regions unparameterized except the largest. The first map is expected to be the least noisy map, but with a potential to hide currently unmodeled densities. The second map is potentially noisier but completely unbiased with respect to the bulk solvent modeling; this map can be considered as an automatically calculated (with no need to manually specify solvent region to omit) *Polder* map (Liebschner *et al*., 2017).

## 3. Results

### 3.1. Data and models

The mosaic modeling was applied to models and data obtained from the PDB (Burley *et al*., 2021) and we used the same set of entries as in A-13. Most tests were performed with structures in space group P1. This simplifies counting isolated regions, removes the need for checking for symmetry-related copies and so on, which is important for protocol development. Such set is quite representative, consisting of more than 3400 structures of different size and at different data resolutions. During the development stage, several subsets were selected. In the initial tests, only 302 ‘best’ models were considered with low *R*-factor values (below 0.25), solved at a relatively high resolution (*d*_*min*_ ≤ 3.0 Å) and with a high overall completeness (> 95%), and the same for the resolution shell from 8 Å to infinity (also > 95%). Additionally, we excluded both very large models so that tests could be performed quickly and too small models for which calculations on a coarse grid with a step of 1.0 Å was impractical, leaving 227 structures for the initial tests. For advanced tests, to obtain more detailed information using better statistics, all models of the same quality without the size limitation were considered yielding 620 models in total. The final tests were performed with all ∼3400 models. Models with no isolated bulk solvent regions of significant volume, larger than 30 Å^3^, were excluded automatically. Finally, the method was applied a subset of the PDB that was used in A-13, which includes entries in non-P1 space groups. This confirmed the results observed with the structures in P1 (results not shown).

### 3.2. Test description

The aim of the test was two-fold: to exercise the implementation of the new mosaic bulk solvent model on a large set of data and to evaluate its utility compared to the currently used protocol A-13 executed with optimized grid step, solvent and shrink truncation radii (Appendix B). For the global score we used three types of *R*-factors (Booth, 1945) calculated: 1) for the whole data set, 2) using the data of resolution lower than 4 Å (*R*_4_), and 3) using reflections in the lowest resolution shell (*R*_low_). We analyzed also the map values in the bulk solvent regions. All tests were performed independently using two mosaic algorithms, *Algorithm 2* and *Algorithm 4* (see above).

### 3.3. Comparison of algorithms 2 and 4

Unlike algorithm 4, algorithm 2 requires minimization iterations and the calculation of the target and gradients, therefore it is slower than 4. The difference between the scale coefficients *k*_*n*_(**s**) found by the two algorithms is usually within 10-20%, which is reasonable given the very weak contribution of these regions to the overall structure factors. The difference is smaller in low-resolution shells and may increase at higher resolution where the signal approaches noise. The average difference of *R*_4_ between the algorithms calculated across all test cases is 0.04% which is much smaller than what we consider a significant difference (0.25%; Appendix C). In 91% of cases algorithm 4 yields lower *R*_4_ than algorithm 2. Excluding cases where the difference between algorithms 2 and 4 is insignificant (e.g., below 0.25%) results in 100% of cases where algorithm 4 produces a lower *R*_4_.

Summarizing, we observe that algorithm 4 is faster and consistently produces the same or lower *R*_4_ values. We did not see any cases were restraints or constraints on the refined scales *k*_*n*_(**s**) were required. We therefore chose algorithm 4 as the default for the mosaic protocol. However, algorithm 2 would still need to be applied in the case of twinned data (not discussed in this work).

### 3.4. R-factor analysis

For each PDB structure the results of the mosaic model were compared with those obtained with A-13 using an optimized grid step and radii for the mask calculation. For each model we calculated the differences between *R*_*4*_-factors after mosaic and after A-13. Since the size and number of isolated regions is small compared to the whole unit cell, changes in *R*-factors are expected to be rather small. In 8% of cases the *R*-factor difference was insignificant (less than 0.25%; Appendix C), with 97.6% of cases having lower *R*_*4*_ using mosaic model. For the rest of the structures, we observe an overall improvement in *R*_*4*_ by 0.52% with the largest difference being 1.4% (3ejo); *R*_*4*_ was lower using the mosaic model in all cases. As anticipated, the impact on the overall *R* factor is smaller since the mosaic bulk solvent models low-resolution features. The mosaic model reduced the overall *R* in 99.5% of cases with the average improvement of 0.2%. Figure 3 shows the typical *R* factor improvement across the resolution range.

**Figure 3.**
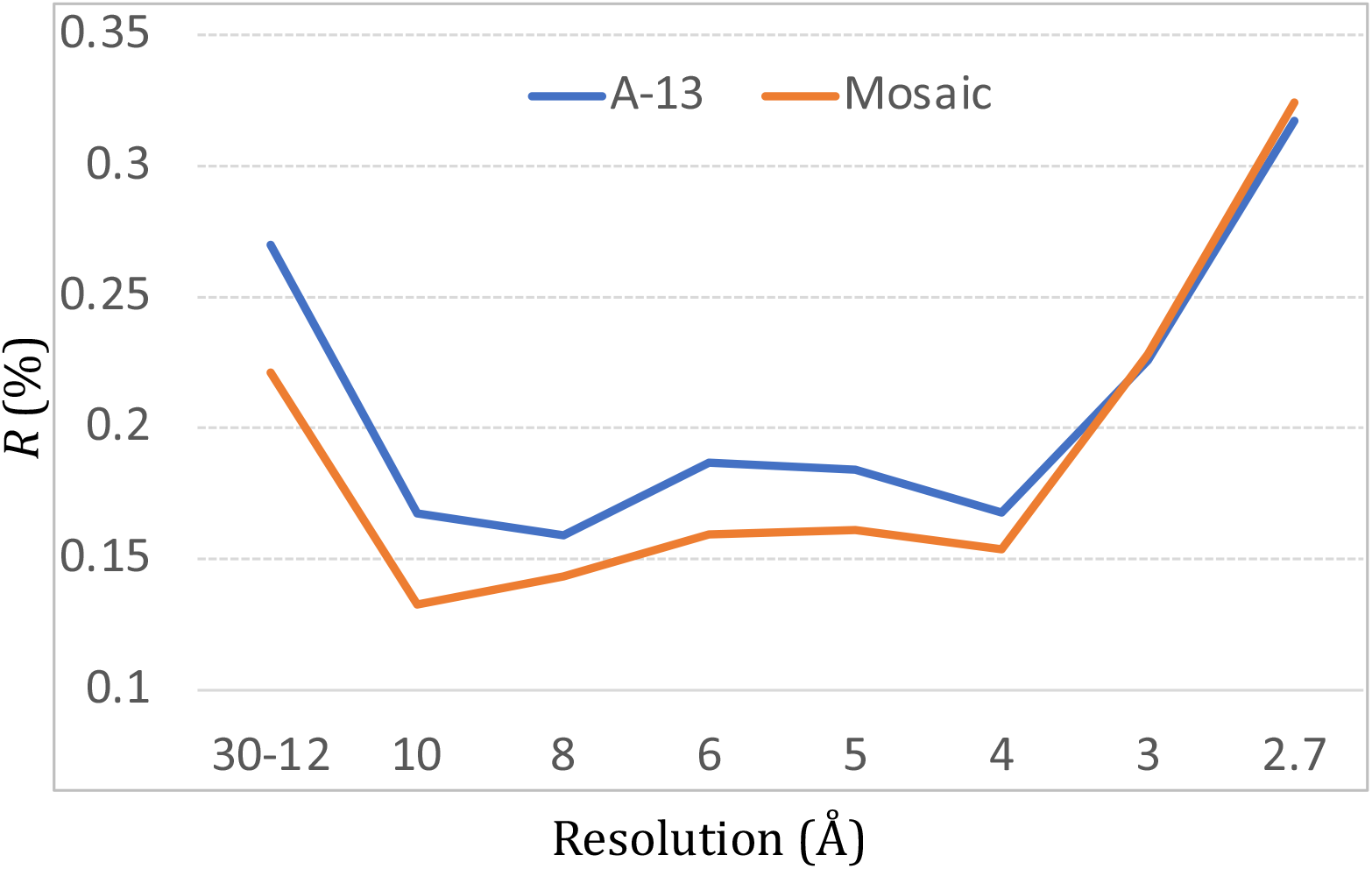
*R* factor per resolution for PDB entry 3e5d shown using A-13 (blue) and mosaic bulk-solvent model (orange).

### 3.5. Map analysis

The results obtained with Algorithm 2 and 4 were very similar, therefore below we present only those for Algorithm 4.

Figure 4 shows the frequency of mean mF_obs_-DF_model_ map (Read, 1986) sigma-normalized electron density values in the isolated regions calculated with the whole mask (the same bulk solvent constant assigned to all solvent regions), with the main mask (bulk density only in the largest component of the solvent region) and after the full mosaic procedure. All maps were normalized to be on the same scale as the map calculated with whole mask. When the whole mask is used there are a substantial number of regions with negative mean density indicating that there is more bulk solvent placed there than needed. In contrast, using the main mask only shows many regions with significant positive density indicating a lack of bulk solvent in these regions. Finally, using the mosaic model eliminates both problems: the maps obtained with the mosaic model contain virtually no regions with mean density beyond ± 0.3 σ.

**Figure 4.**
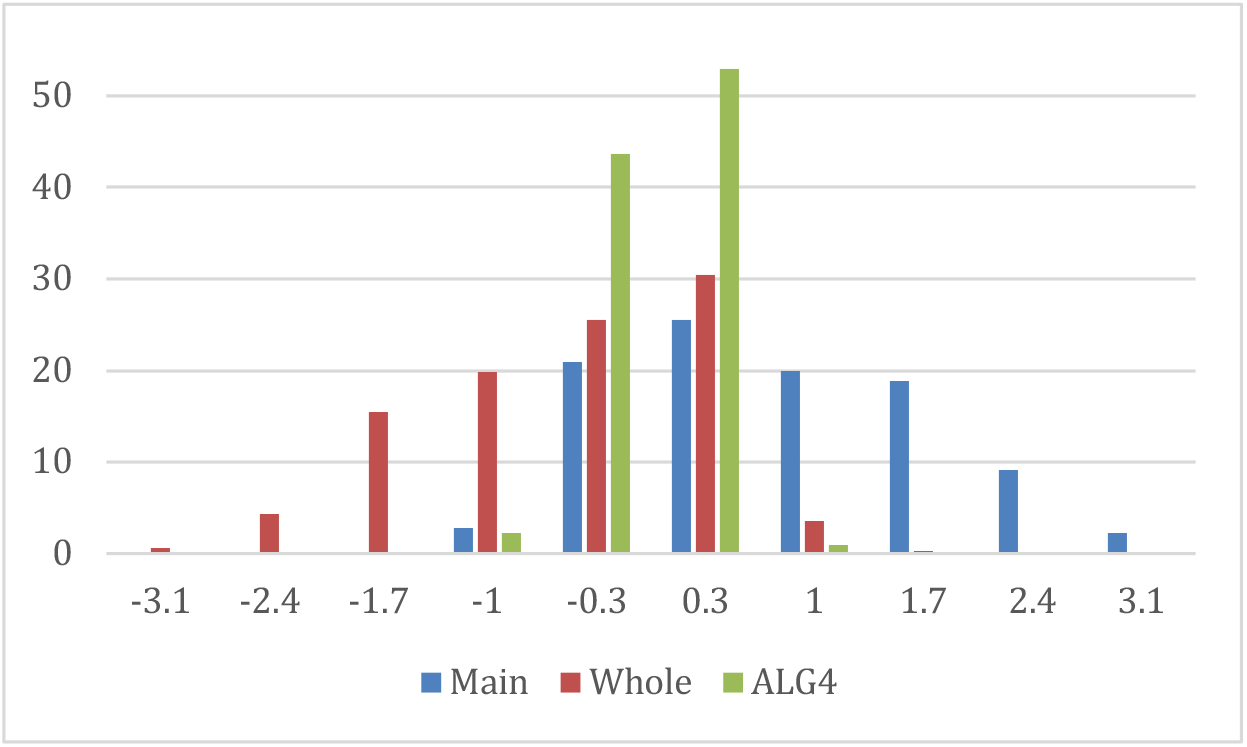
Distribution of mean mF_obs_-DF_model_ map values calculated in isolated bulk solvent regions, all scaled in r.m.s. of the map calculated with the ‘whole’ mask.

Figure 5 shows three typical scenarios for the regions: an empty region (left), a region filled with a different solvent than elsewhere (middle) and region filled with disordered solvent (right). Indeed, applying A-13 with the solvent mask calculated for the largest main region only shows no density in the empty region and large residual positive peaks in the other two regions (Fig. 5b). Modelling the solvent with A-13 using the default (whole) mask results in negative difference map artifacts in the empty region (Fig. 5c, left), it fully models the disordered solvent (Fig. 5c, right) but it only partially accounts for the solvent in the other region (Fig. 5c, middle). Finally, applying mosaic model successfully treats the solvent occupation (or lack thereof) in all three regions (Fig. 5d).

**Figure 5.**
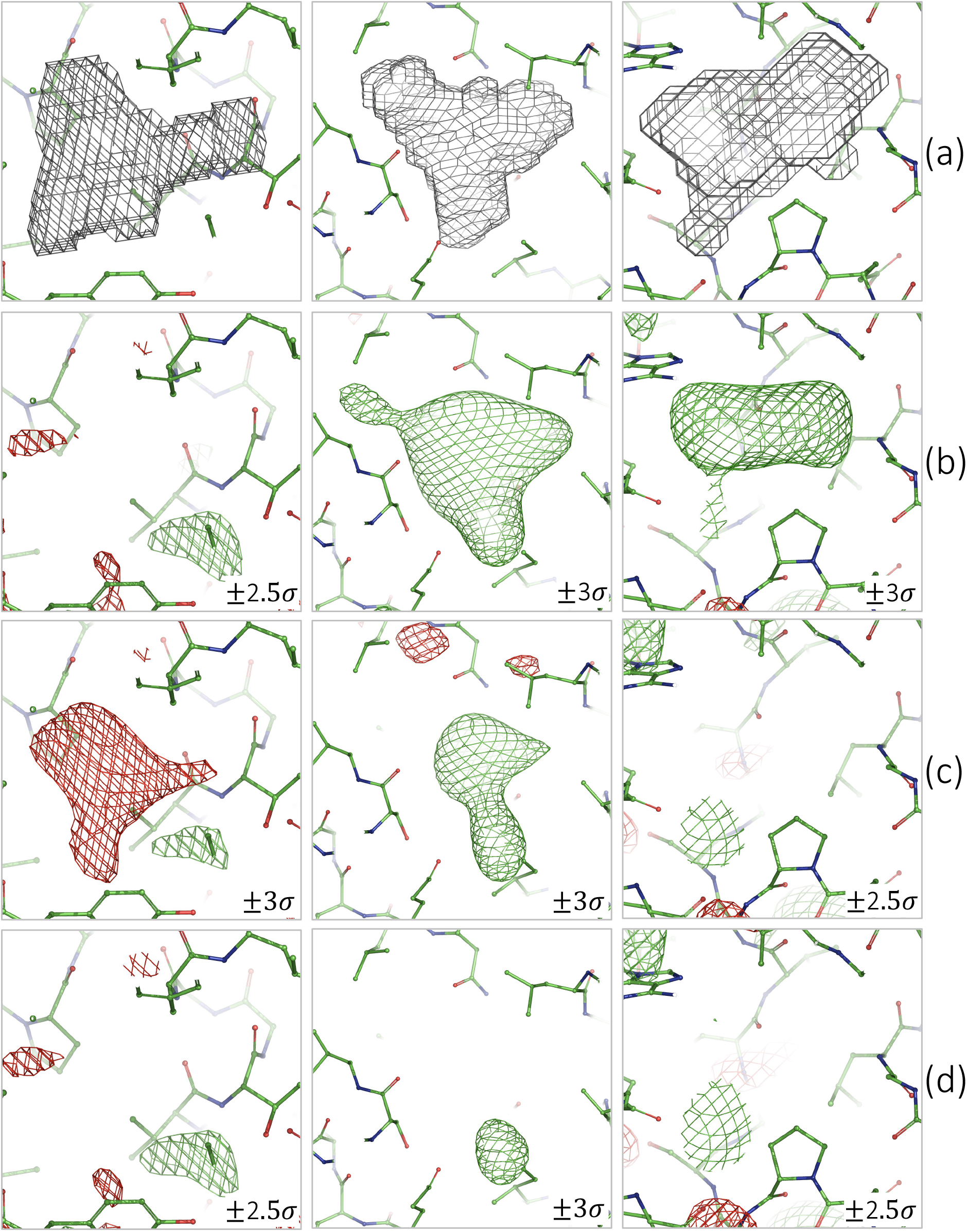
Examples of isolated regions: near residue 809 in chain D (PDB: 4guo; left), near residue 131 in chain H (middle) and near residue 271 in chain A (right) for 3blw. (a) Bulk-solvent mask. mF_obs_-DF_model_ maps with F_model_ calculated: (b) using one largest bulk-solvent mask, (c) as in A-13 and (d) using mosaic bulk solvent model. Contouring levels are indicated on pictures. Positive and negative contours are in green and red, respectively.

### 3.6. Simplified mosaic protocol

Using the simplified mosaic model with algorithm 4 (2.1.3) slightly increases *R*_*4*_ compared to the full version in 92.7% of cases. However, the average *R*_*4*_ difference between the simplified and full versions is 0.02% for cases where *R*_*4*_ increased and 0.06% for cases where *R*_*4*_ decreased; both of which are below the significance level of 0.25%. Map analysis shows virtually the same behavior of both protocols for this subset of structures. A larger difference is anticipated for structures that include a mixture of different solvent types, e.g., semi-ordered lipids in membrane proteins.

## 4. Conclusions

The mosaic bulk solvent model addresses a number of drawbacks of the flat mask-based solvent model (JB-94; A-13). This approach makes it possible to model solvent regions independently and in a computationally efficient manner, in turn allowing for partial solvent occupancy in these regions and potentially different kinds of solvent regions, such as lipids. Using the mosaic solvent model improves the overall fit of the model to the data and reduces artifacts in residual maps.

The mosaic model algorithm was systematically exercised against a large subset of PDB entries to ensure its robustness, computational efficacy and practical utility to improve maps and models compared to the current protocol (A-13). While the mosaic protocol is by design more computationally expensive than A-13, the benefits of using it are likely to outweigh the computational cost in most cases. As we demonstrate in our tests, using the mosaic model systematically improves low-resolution *R*-factors by about 0.5% and removes residual map artifacts due to insufficiency of the flat model. The mosaic solvent approach can also be extended in the future to allow for more efficient modeling of partially disordered solvent with unique characteristics, such as lipids belts in membrane proteins. We plan to investigate this topic as a follow up work.

The mosaic solvent model is intended to account for finer details that become important towards the end of model refinement. Therefore, it best applied at the later stages of structure solution, and not in initial stages when gross modeling errors are still to be corrected. It is similar to using explicit hydrogen atoms in atomic models. Using hydrogen atoms at initial stages is likely to be counterproductive as they may hinder the correction of larger-scale modeling errors, while using them towards the end can improve *R* factors by about 1% (Afonine *et al*., 2010; Afonine & Adams, 2012) and local geometry (Williams *et al*., 2018).

The combination of optimal parameters for the mask calculation (grid step, solvent and shrinkage radii) found as (0.6, 1.1 and 0.9 Å) is universal and not specific to mosaic solvent modeling approach.

The algorithms described here are implemented in CCTBX and are used in the *Phenix* suite (Liebschner *et al*., 2019) where applicable starting from version 1.20rc4-4425.

## Acknowledgments

PVA and PDA thank the NIH (grants R01GM071939, P01GM063210 and R24GM141254) and the PHENIX Industrial Consortium for support of the PHENIX project. This work was supported in part by the US Department of Energy under Contract No. DE-AC02-05CH11231. AU thanks the French Infrastructure for Integrated Structural Biology (FRISBI) ANR-10-INSB-05-01 and Instruct, which is part of the European Strategy Forum on Research Infrastructures (ESFRI) and is supported by national member subscriptions.

All map and model figures were prepared with *Pymol* (DeLano, 2002).

## Appendix A Composition of the solvent mask

Crystallographic modeling assumes any voxel of the unit cell that is not occupied by the atomic model to be occupied by the disorder solvent. The bulk-solvent region may not be just a large region between macromolecules but can also form local cavities inside macromolecular region or on the interface between the macromolecular chains inside the asymmetric unit or across domains of the same or different molecular copies. In vast majority of cases bulk solvent regions consist of one large region (which we refer to as ‘main’ region) and some number of much smaller secondary regions. Table A1 shows the distribution of such secondary components with a volume larger than 30, 50, 100 or 200 Å^3^ for a set of 3237 models in space group P1. As the extreme case, the structure 2glj has 475 regions of the volume larger than 30 Å^3^. For most of these structures, the second largest region has a volume much smaller that the main one (Fig. A1).

## Appendix B Choice of the mask parameters

Bulk solvent mask calculation requires four parameters to be defined: mask grid step (*d*_*grid*_), solvent (*r*_*solv*_) and shrinkage (*r*_*shrink*_) radii, and atomic radii that are chemical element-specific. While the latter are the tabulated values centrally available to major software packages, the choice of the other three is less straightforward, implementation-specific and may be a subject of optimization. Assuming these mask calculation parameters are independent of the data set resolution, the aim here is to find a universal set of these parameters that are applicable to any atomic model. The major difficulty is an expected high correlation of *d*_*grid*_ with the *r*_*solv*_ and *r*_*shrink*_. To search for the universal optimal values of these parameters, we started the analysis from 302 quality-filtered models as described in § 3.1. The *R*-factor calculated in the resolution interval from infinity to 4 Å (*R*_4_) was used here to evaluate parameter choices.

The optimal parameter search was based on a three-dimensional systematic search over a range of *d*_*grid*_ (from 0.2 to 1.0), and *r*_*solv*_ and *r*_*shrink*_ ranging from 0 to 1.6 Å. For each triplet values of trial (*d*_*grid*_, *r*_*solv*_, *r*_*shrink*_) the A-13 scaling protocol was applied and the value of *R*_4_ was calculated for each of the models. We refer to *R*_*opt*_(*d*_*grid*_) as the lowest value found for a given structure and a given grid step over all scanned *r*_*solv*_ and *r*_*shrink*_ values.

First, we observed that *R*_*opt*_(*d*_*grid*_) varies similarly for different structures. It practically does not change for *d*_*grid*_ in the range 0.2 – 0.4 Å (meaning that such step is unnecessarily small) while increases sharply start at around 0.7 – 0.8 Å (meaning that such step becomes too coarse). Fig. B1a shows the average curve of such variation.

Second, this observation is true also for two other types of masks: when all isolated connected regions of the volume below 30 Å^3^ were excluded from the mask, and also when all regions except the largest one (‘main’) were excluded. Moreover, *R*_*opt*_(*d*_*grid*_) values for these masks are systematically lower than those for the initial mask (the ‘whole mask’). At the same time, the *rmsd* values (Fig. B1b) indicate the presence of exceptions. For the masks calculated with the hydrogen atoms included into the model, *R*_*opt*_ was systematically larger than without hydrogens and later we did not consider such masks any more.

Third, using the selected above range of *d*_*grid*_, we noted the preferred values for *r*_*solv*_ and *r*_*shrink*_, those for which *R*_*opt*_(*d*_*grid*_) is obtained. To keep track of eventually possible several choices, we calculated a kind of a Boltzmann-energy function maxima of which indicate the preferable (*r*_*solv*_, *r*_*shrink*_) combinations for each *d*_*grid*_. To do so, for each set of parameters (*r*_*solv*_, *r*_*shrink*_; *d*_*grid*_) and for each structure *n* we calculated a value

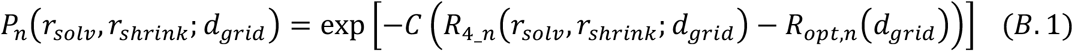

and then a product of such values over all structures:

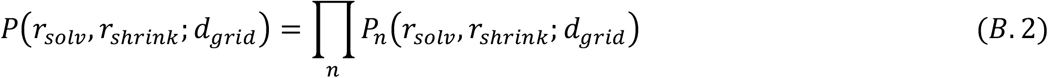

For the given set of structures, an appropriate value for the constant *C* found empirically was equal to 1. Distributions (B.2) indicated a clear correlation between *r*_*solv*_ and *r*_*shrink*_ and a relative pertinence of the optimal values (not shown). For the grid steps we are interested in, 0.5-0.7 Å, the optimal radii (*r*_*solv*_, *r*_*shrink*_) varied around the values (1.1 Å, 0.9 Å) while we did not discard several neighboring combinations.

Finally, we checked how much *R*_4_(*r*_*solv*_, *r*_*shrink*_, *d*_*grid*_) obtained with these chosen triplets of values (*r*_*solv*_, *r*_*shrink*_, *d*_*grid*_) is worse than *R*_4_*best*_, the value obtained by optimization of all three parameters for each model. To do so, we calculated for how many test structures the difference

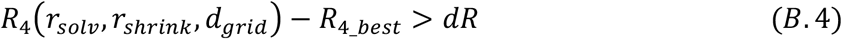

is above different threshold values *dR*. Fig. B2 gives several examples. The best would be the curve approaching the horizontal axis at the smallest threshold and going as low as possible for *d*_*grid*_ = 0.6 Å and eventually at 0.5 Å or 0.7 Å. All curves on the left plot (*r*_*solv*_ = 0.9 Å, *r*_*shrink*_ = 0.7 Å) are higher than the curves on the central plot (*r*_*solv*_ = 1.1 Å, *r*_*shrink*_ = 0.9 Å) and by this reason we reject the first combination. On the plot on the right, two curves are closer to the horizontal axis but they correspond to too small steps that we excluded. At the same time, the curves corresponding to the step we are considering are also higher than the curves on the central image. Further analysis of the curves and respective tables confirmed the best choice for *r*_*solv*_ = 1.1 Å, *r*_*shrink*_ = 0.9 Å, *d*_*grid*_ = 0.6 Å.

## Appendix C Mask filtering

Isolated solvent regions vary by their volume and content. An estimate for the volume of the smallest region that one can still assume to contain solvent is 27 Å^3^ (the volume of a single water molecule). Smaller regions can safely be considered as geometrical artifacts of mask calculation procedure. Remaining regions may still be empty but this cannot be reliably inferred using geometric criteria only, thus extra information is needed. It can be obtained from mF_obs_-DF_model_ difference OMIT map with F_model_ calculated using the main mask only as described in §2.2.

To select the appropriate value for this threshold, first we removed all regions of the volume *V* smaller than 30 Å^3^. Removing this obvious noise on average decreases *R*_4_ except for a few models for which it increased by not more than 0.24% (for a single model it was 0.28%). We consider this value as the margin of an acceptable fluctuation in *R*_4_. Removing regions with *V* < 50 Å^3^ kept *R*_4_ at the same level while a selection by *V* < 80 Å^3^ increased it (Fig. C1). *R*_4_ decreased in comparison with that for the whole mask, *R*_4_whole_, approximately for about 550 models from 620 used in this test (see § 3.1) for each of the three volume threshold values (30, 50, 80 Å^3^), proving that most of removed regions are indeed empty or noisy.

Using different OMIT map one can calculate the minimal, *ρ*_*min*_, maximal, *ρ*_*max*_, and mean, *ρ*_*mean*_, map values within each region. This was done with the same subset of 620 quality filtered models and we observed a strong correlation of *ρ*_*min*_ and *ρ*_*max*_ with *ρ*_*mean*_. Therefore, we left only *ρ*_*mean*_ as an independent characteristic.

After elimination of all small-volume regions with the chosen threshold, we additionally removed the regions with *ρ*_*mean*_ below 0.0, or 0.5, or 1.0 σ. For all combinations of these cut-off values, eliminations with *V* < 80 Å^3^ or *ρ*_*mean*_ < 1.0 σ systematically gave larger *R*_4_ for some structures. For other combinations, the results were similar (not shown) while the combined condition *V* < 50 Å^3^ or *ρ*_*mean*_ < 0.5 σ eliminated more regions, than the others. Thus we selected this condition as the default one for eliminating inappropriate regions.

---

**Figure A1.**
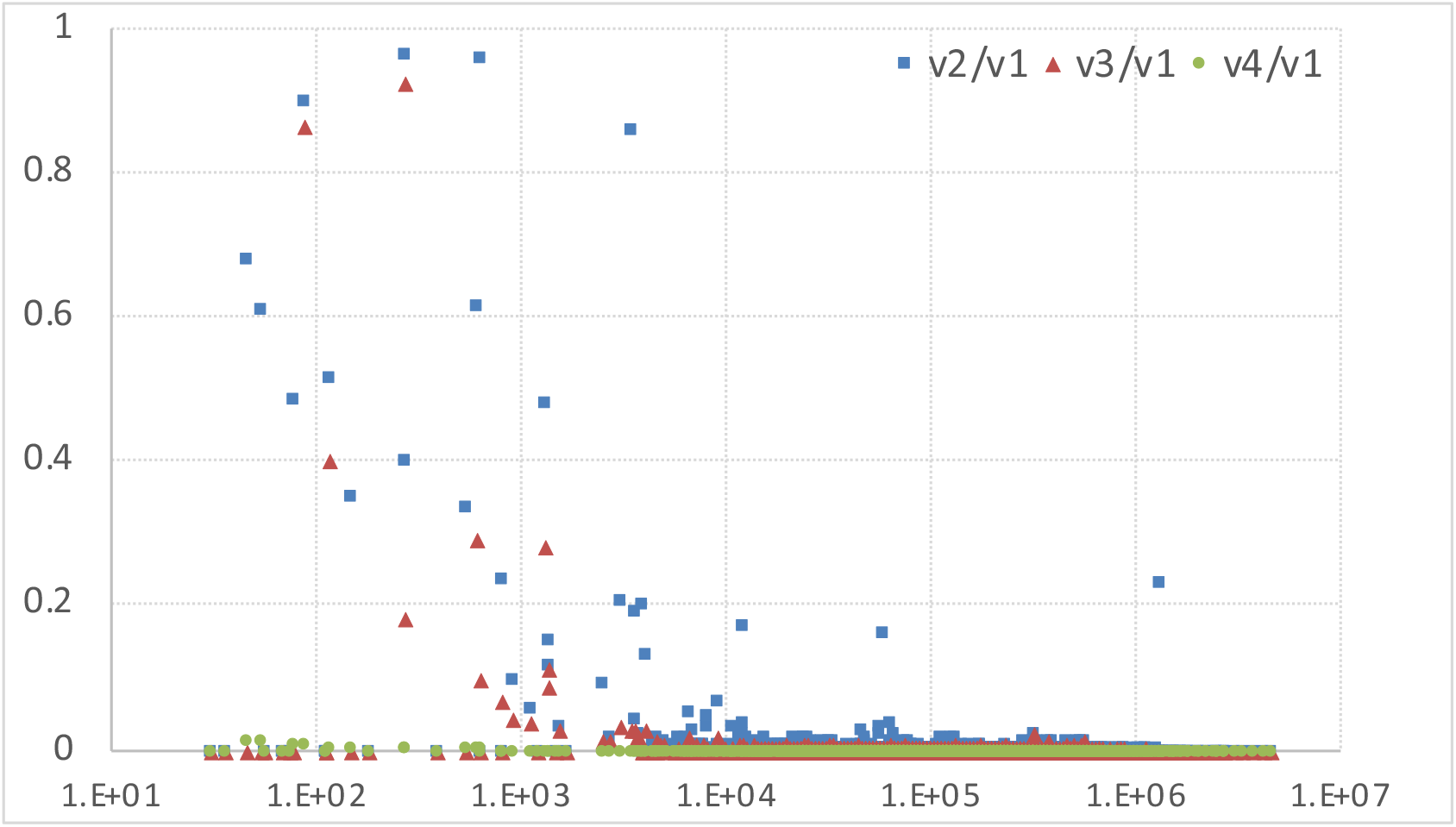
Ratio of the volume of the bulk solvent components to the volume of the main component. Each structure from the test set is represented by three points, all with the abscise showing the volume, in Å^3^, of the first, main component (components are given in decreasing order of their volume). Ordinates of the blue squares, red triangles and green dots are equal to the ratio of the volume of the second, third and fourth components, respectively, to the volume of the first component.

**Figure B1.**
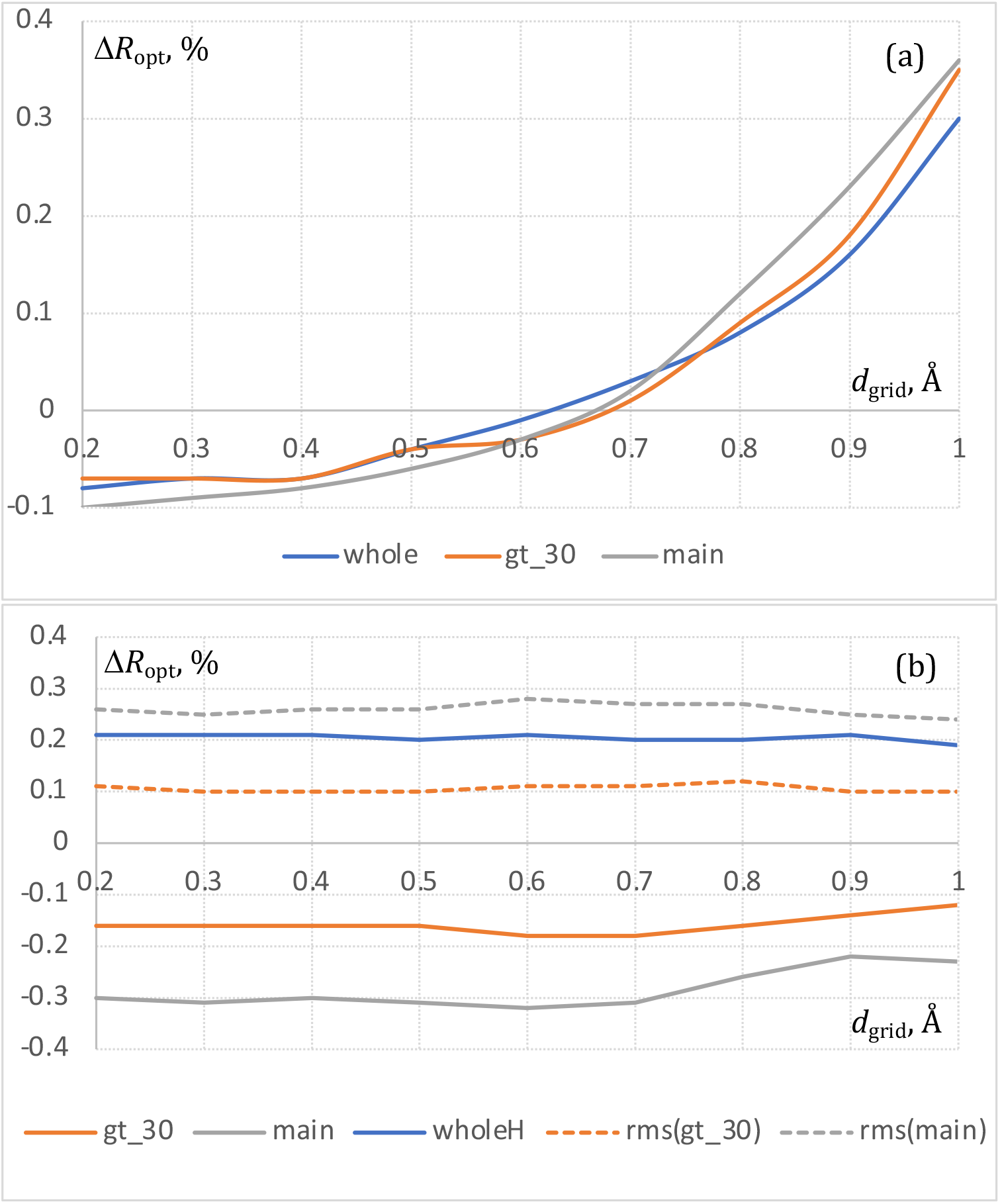
a) Variation of *R*_*opt*_ in comparison with the respective mean value over all steps. Distribution shown for the whole mask, for the mask without small-volume regions (gt_30) and for that composed of the only major component (main). b) Comparison of average *R*_*opt*_ for different masks with the average for the whole mask build without hydrogens.

**Figure B2.**
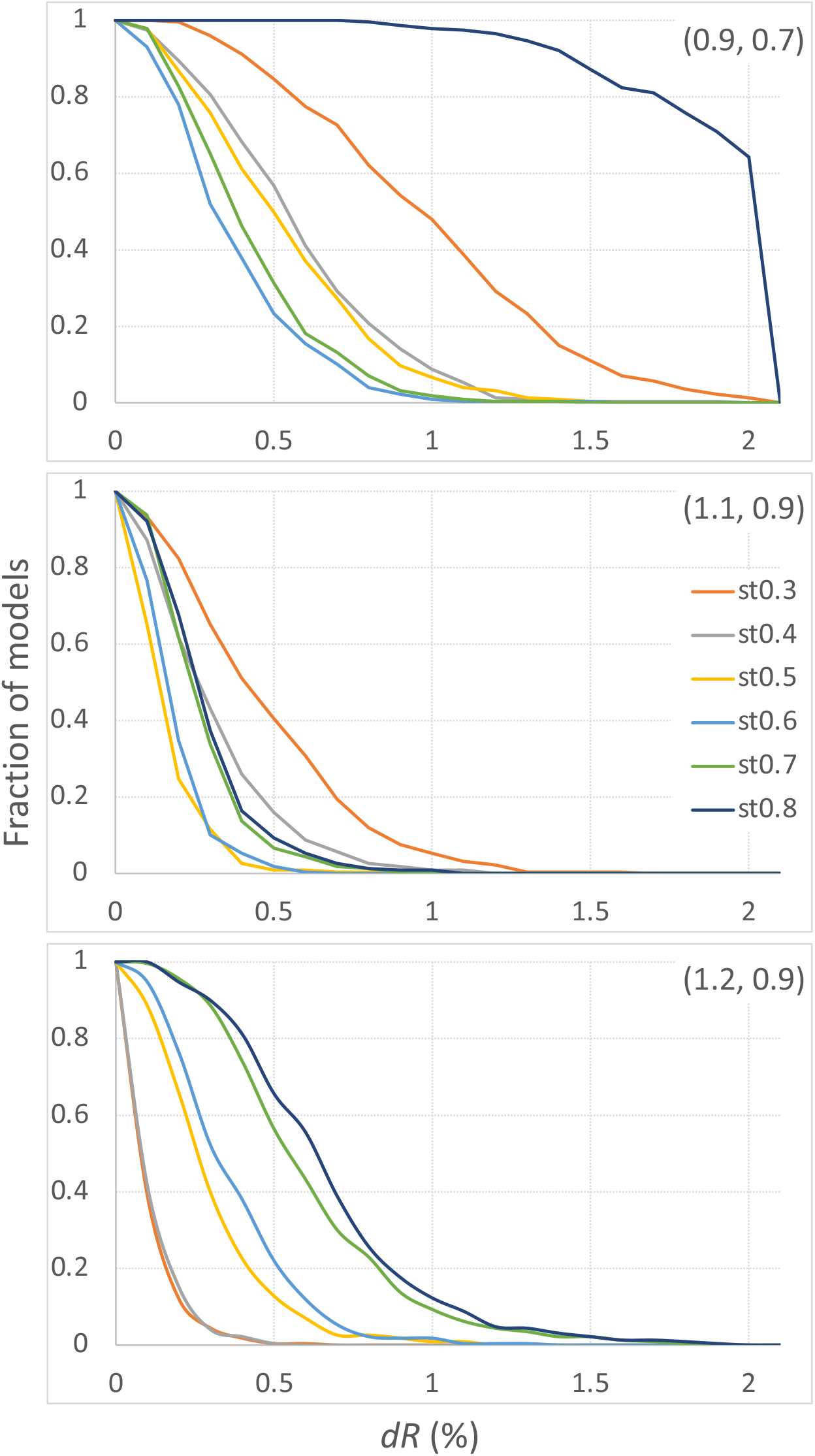
A fraction of models such that, using the given radii (*r*_*solv*_; *r*_*shrink*_) and a different grid step in the range 0.3 – 0.8 Å, *R*_4_ is worse than the best possible *R*_4_ by more than *dR*. Note an outlier curve for *d*_*grid*_ > *r*_*shrink*_ (left) and optimal curves for (*r*_*solv*_, *r*_*shrink*_, *d*_*grid*_) equal to (1.1, 0.9, 0.6) (central plot) or to (1.2, 0.9, 0.4) (right plot).

**Figure C1.**
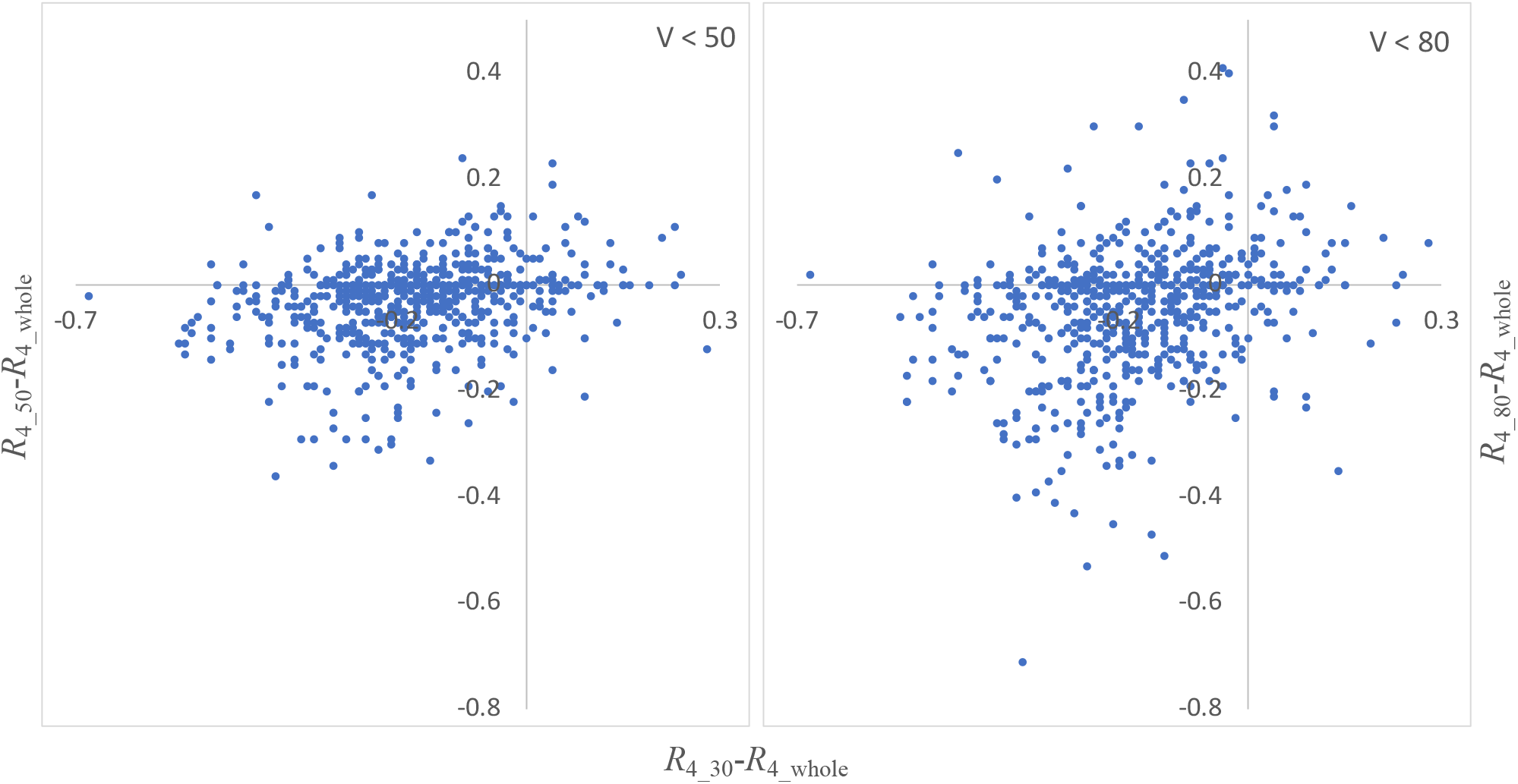
Distributions of *R*-factor differences (in %): *R*_4_30_-*R*_4_whole_ (both plots) versus *R*_4_50_-*R*_4_whole_ (left) and *R*_4_80_-*R*_4_whole_ (right). Dots in the left figure are closer to the horizontal axis showing that removing the regions of the volume between 30 and 50 Å^3^ improves *R*_4_ further, which is not the case when removing the regions of the volume between 30 and 80 Å^3^ (right figure).

**Table A1.** Number of isolated components of the bulk solvent region. For each volume cut-off *V*, in Å^3^ (left column), the table shows the percentage of model that have no components of the volume larger than *V* (column “0”), have one such component (column “1”), two such components (column “2”), and so on. Column “Nmax” shows, for the components of the volume larger than *V*, the maximal number of such components per structure observed in the test set.

| Volume | Nmax | 0 | 1 | 2 | 3 | 4 | 5 | 6 | 7 | 8 | 9 | 10 | >10 |
| --- | --- | --- | --- | --- | --- | --- | --- | --- | --- | --- | --- | --- | --- |
| 30 | 475 | 0.17 | 8.32 | 7.97 | 7.77 | 7.36 | 6.98 | 6.55 | 5.32 | 5.03 | 4.04 | 3.93 | 36.54 |
| 50 | 289 | 0.26 | 23.39 | 15.59 | 11.84 | 8.32 | 7.30 | 5.32 | 4.51 | 3.11 | 2.44 | 1.75 | 16.15 |
| 80 | 163 | 0.41 | 46.67 | 17.66 | 10.71 | 5.44 | 4.19 | 3.00 | 2.44 | 1.45 | 1.25 | 0.70 | 6.08 |
| 100 | 112 | 0.44 | 58.02 | 15.68 | 9.08 | 4.25 | 3.38 | 2.15 | 1.28 | 0.90 | 0.70 | 0.47 | 3.67 |
| 120 | 93 | 0.52 | 66.02 | 13.50 | 7.39 | 3.61 | 2.53 | 1.60 | 0.99 | 0.84 | 0.41 | 0.41 | 2.18 |
| 150 | 48 | 0.52 | 74.51 | 11.03 | 5.96 | 2.39 | 1.77 | 1.22 | 0.49 | 0.26 | 0.32 | 0.29 | 1.22 |
| 200 | 28 | 0.58 | 82.72 | 8.84 | 3.64 | 1.45 | 1.11 | 0.55 | 0.15 | 0.29 | 0.17 | 0.09 | 0.41 |
| 250 | 21 | 0.58 | 87.93 | 5.99 | 3.05 | 0.81 | 0.81 | 0.20 | 0.15 | 0.06 | 0.09 | 0.06 | 0.26 |
| 300 | 12 | 0.64 | 90.69 | 4.68 | 2.27 | 0.67 | 0.49 | 0.20 | 0.06 | 0.03 | 0.17 | 0.03 | 0.06 |
| 400 | 11 | 0.67 | 93.37 | 3.17 | 1.95 | 0.49 | 0.23 | 0.03 | 0.03 | 0.00 | 0.03 | 0.00 | 0.03 |
| 500 | 9 | 0.67 | 95.00 | 2.09 | 1.75 | 0.26 | 0.17 | 0.03 | 0.00 | 0.00 | 0.03 | 0.00 | 0.00 |

